# Evidence of shifts towards neural states of stability during the retrieval of real-life episodic memories

**DOI:** 10.1101/2020.06.12.147942

**Authors:** L Fuentemilla, B Nicolás, A Kastrinogiannis, M Silva

**Author notes:** **Contact information.** Lluís Fuentemilla, Department of Cognition, Development and Educational Psychology, University of Barcelona. Pg Vall Hebrón, 171, 08035 Barcelona, Spain.

## Abstract

How does one retrieve real-life episodic memories? Here, we tested the hypothesis, derived from computational models, that successful retrieval relies on neural dynamics patterns that rapidly shift towards stable states. We implemented cross-temporal correlation analysis of electroencephalographic (EEG) recordings while participants retrieved episodic memories cued by pictures collected with a wearable camera depicting real-life episodes taking place at “home” and at “the office”. We found that the retrieval of real-life episodic memories is supported by rapid shift towards brain states of stable activity, that the degree of neural stability is associated with the participants’ ability to recollect the episodic content cued by the picture, and that each individual elicits stable EEG patterns that were not shared with other participants. These results indicate that the retrieval of autobiographical memory episodes is supported by rapid shifts of neural activity towards stable states.

## Introduction

The retrieval of episodic memories is thought to rely on neural mechanisms that allow rapid association of multiple event elements from one’s past experience to forge a self-consistent memory representation (Tulving, 2002; Eichenbaum et al., 2007). Theoretical models of memory agree that this is supported by a dynamic interaction of the hippocampus and neocortical regions regulated by attractor-like dynamics (Marr, 1971; McClelland et al., 1995; McNaughton and Morris, 1987; Norman and O’Reilly, 1994), which refers to the property of the neural network to shift towards a stable state when the external inputs are correlated, but not identical, to a stored pattern (Hopfield, 1982; Treves and Rolls, 1997). Indeed, this is a necessary requirement for retrieving memories from everyday life activity, as they are often contextually overlapping but unique in their episodic content. However, while attractor-like network dynamics in the human brain, such as shifts towards stable states, have been reported in perceptual categorization (Quian-Quiroga et al., 2014; Rotshtein et al., 2005), in working memory (Kamiński et al., 2017) and in spatial navigation (Steemers et al., 2016) it remains elusive in the retrieval of episodic memories.

Based on extensive electrophysiological (EEG) evidence showing that mnemonic processes can take place rapidly in humans, i.e., within the first 1500 ms from a reminder onset (Yonelinas, 2002; Rugg and Curran, 2007; Staresina and Wimber, 2019), here we tested the hypothesis that neural activity following a retrieval cue would settle into a stable state within 1500 ms after a cue onset. Importantly, and in line with the theoretical prediction that cue-eliciting neural network states of stability are content-specific in memory (Rolls, 2016), we hypothesized that memory cues depicting event episodes experienced under similar real-life category contexts, such as those that take place “in my home” or that happen “in my office”, would entail similar neural states of stability. Indeed, arguably, the use of multivariate pattern analyses in experimental research using pictures and words encoded in the lab (Norman et al., 2006; Kriegeskorte et al., 2008) has furnished evidence for neural reinstatement of category features at retrieval (Polyn et al., 2005; Johnson and Rugg, 2007; Jafarpour et al., 2014; Horner et al., 2015; Ritchey et al., 2013; Staresina et al., 2012; Tompary et al., 2016). However, this research strategy comes at a cost, as it cannot capture an essential feature of everyday life episodic memory functioning, which is that episodes are experienced at individual bases, and their retrieval should, consequently, be mediated by idiosyncratic neural patterns of activity thereof.

To test these hypotheses in ecologically-valid scenarios, we asked participants to use a wearable camera to automatically capture pictures depicting real-life episodes taking place at their “home” and at their “office”, and we then used these pictures to cue the retrieval of these past episodes in the lab, while scalp EEG was recorded. We show that pictures depicting past episodes elicited a neural response activity that shifted towards a stable state after ~ 800 ms from picture onset. Importantly, we found that a similar stable neural pattern was elicited by pictures depicting experienced episodes, but not novel ones, in the same context (i.e., at home or at the office), and that this pattern of stable brain activity was not shared across individuals. Finally, we replicated these findings in data from a second experiment with different participants and extended them by showing that the degree of context-specific neural stability is associated with the participants’ ability to recall contextual information from the cued episodes.

## Results

### Assessing memory for real-life episodic events

Participants were instructed to carry a wearable camera during their routine daily activity over 5-7 consecutive days; the camera took face-front pictures automatically every 30 seconds. Many of the participants’ sets of recorded pictures depicted episodic instances taking place at their “home” or at their working “office” (Fig. 1a). Thus, in the current study, we focussed on analysing participants’ responses to pictures from these two real-life contextual categories. Furthermore, pictures from these two real-life contexts included elements and features that had much more in common between these two categories (they depict indoor scenes, they show similar furniture, etc.) than with other categories, such as events happening “on the street” or “on public transport”. This minimized the possibility that neural response differences between context categories could be explained simply by general layout properties of the pictures. After encoding, a selection of these pictures was used in the lab to cue participants’ memories of specific event episodes, while EEG was recorded.

**Figure 1.**
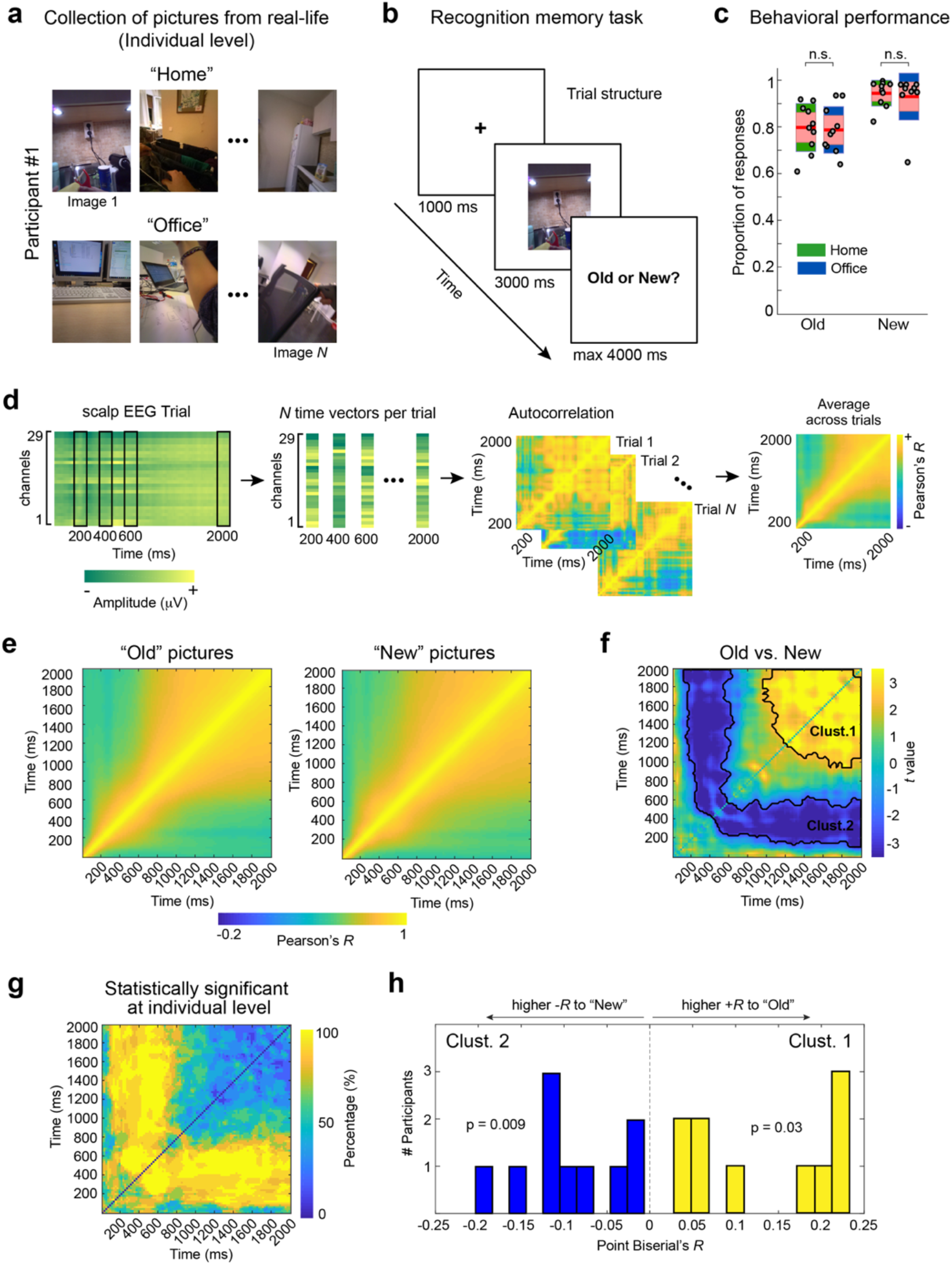
Task design and results in experiment 1. (**a**) Example of pictures collected by the wearable camera at “home” or at “the office” by one of the participants in experiment 1. (**b**) Example of a trial in the recognition memory test, where each individual had to indicate whether the picture presented belonged to her past (Old) or not (New). (**c**) Participants’ proportion of correct responses to ‘Old’ and ‘New’ pictures as a function of the depicted context. The central mark is the median, the edges of the box are the 25th and 75th percentiles. ‘n.s.’ denotes p > 0.05. (**d**) Schematic representation of the analysis. A temporal autocorrelation matrix (Pearson’s *R)* is generated from raw EEG data for each of the participants. Patterns of neural response to each time point after cue onset correlated to the rest of the neural patterns throughout 2 seconds from cue onset for each trial separately are then averaged. (**e**) Correlation matrices display the temporally resolved correlation patterns for separately averaged single trial 2D correlation maps for EEG response to Old and New pictures across subjects. (**f**) 2D map of t-values resulting from comparing Old and New correlation maps across participants. The thick black line depicts the timing of the significant cluster between conditions (p < 0.05, cluster-based permutation test). (**g**) Point-to-point percentage of participants that showed statistical significance (p < 0.05) at individual level when comparing 2D correlation maps from Old and New pictures. (**h**) Histogram of the participants’ point biserial correlation coefficients separately for each of the two clusters identified in (f). p values resulting from testing (Student t-test) whether the group regression estimates differed statistically from 0.

In experiment 1, a recognition memory task was implemented to assess participants’ (*N* = 10) memory of past episodes one and two weeks after encoding. In this task, a selected set of pictures depicting events taking place at “home” or at “the office” (Old pictures) were presented interspersed with pictures form the same contextual categories recorded by another participants (New pictures). Participants were asked to indicate whether the picture presented on the screen depicted episodes from their past or not (Fig. 1b). Overall, we found that participants were highly accurate in identifying Old and New pictures irrespective of whether they depicted home or office scene episodes (repeated measures ANOVA; main effect of picture category: F(1,9) = 0.65, p = 0.44; interaction picture category x response type: F(1,9) = 0.16, p = 0.90) (Fig. 1c).

### Temporally evolving neural states of stability

To examine whether EEG response patterns elicited by Old and New pictures evolved towards stable states, we implemented a cross-temporal autocorrelation approach (Fig. 1d). Briefly, a time-to-time Pearson correlation analysis to each individual’s trials quantified the degree of similarity between a neural pattern at a given time point and the rest of the neural patterns elicited by a picture throughout a time window of 2 seconds from picture onset. The extent to which a neural pattern is stable in time should be seen as an increase in correlation values surrounding that time point. This analysis revealed that EEG response patterns to Old and New pictures gradually generalize over time (Fig. 1e). However, we found that the degree of pattern stability induced by New pictures was greater in early time windows after picture onset whereas Old pictures induced a more stable pattern of neural activity at later stages, at ~ 800 ms after picture onset (Fig. 1f). Therefore, in line with our hypothesis, pictures cueing the retrieval of episodic memories induced shifts towards stable states of activity, which are dissociated from those elicited by pictures that cued no memory content.

Next, we sought to investigate whether the dissociated states of stability for Old and New picture cues was consistent at the individual level. To address this issue, we implemented a time-to-time t-test comparison across trials for each individual and calculated the percentage of participants that showed a significant (p < 0.05) effect. This analysis revealed that the time-resolved EEG correlation cluster effects seen at the group level were highly consistent at the individual level (Fig. 1g). To assess these effects statistically, we performed a Point-Biserial regression analysis at the individual level using a correlation value extracted from averaged neural similarity of sample points included in clusters 1 and 2 identified in the group level analysis (i.e., Fig. 1f) as dependent variable and picture category (Old or New) as independent measure. If neural similarity for each of the clusters scaled according to picture condition, then the slope of this relationship should differ from 0 and in the opposite direction for each cluster. Indeed, this was the case, as cluster slopes were positive for Cluster 1 and negative for Cluster 2, and the two of them were significantly different from 0 (Cluster 1: t(9) = 3.91, p = 0.03; Cluster 2: t(9) = −4.79, p = 0.009) (Fig. 1h). Altogether, these findings suggest that the time course of neural activity during retrieval shifted towards neural states of stability at different temporal windows as a function of whether external inputs matched or mismatched internal memory representations.

### Shift towards neural stability in episodic memory retrieval

We next examined whether EEG states of stability were associated with accessing real-life contextual category information depicted by the picture. To address this issue, we implemented a cross-temporal correlation analysis whereby half of the EEG trials from one contextual category were correlated to the other half of the same (Within Category condition; for example, “home” with “home”) or to EEG trails from the other category (Between Category condition; for example, “home” with “office”). To assess the retrieval-dependent nature of the effects, this analysis was implemented with Old and New trials in like manner. The results of this analysis revealed significantly greater neural stability from ~1000 ms after picture onset only when the similarity analysis included trials within the same contextual category in Old responses (Fig. 2 and Supplementary Fig. 1). These results were corroborated after running the same analysis on similarity values extracted from randomly assigning set of trials into 2 halves 1000 times (Supplementary Fig. 2), thereby ruling out the possibility that the increase in neural stability found in the main analysis in the Within condition compared to the Between condition was artificially induced by how EEG trials were split into two sets before running the correlation analysis. Finally, a repeated measures ANOVA with correlation values from the cluster confirmed that the increased neural similarity in the Within as compared to the Between condition was specific to Old trials, as it revealed a significant Response Type (Old and New) x Content type (within and between) interaction effect (F(1,9) = 7.47, p = 0.02) and a significant main Response Type effect (F(1,9) = 7.27, p = 0.02) but not a Content type effect (F(1,9) = 0.06, p = 0.81). A paired *t*-test confirmed the ANOVA interaction was driven by differences in the Within vs Between contrast for Old responses (t(9) = 2.48, p = 0.03) but not for New responses (t(9) = −1.25, p = 0.24) (Supplementary Fig. 3). Altogether, these results suggest that shifts towards neural stability observed after ~800 ms from stimulus onset were associated with retrieving episodic memories and that the pattern of stability was specifically associated with the memory content. These findings lend support to the theoretical prediction that cue-eliciting neural network states of stability are content-specific in memory (Rolls, 2016).

**Figure 2.**
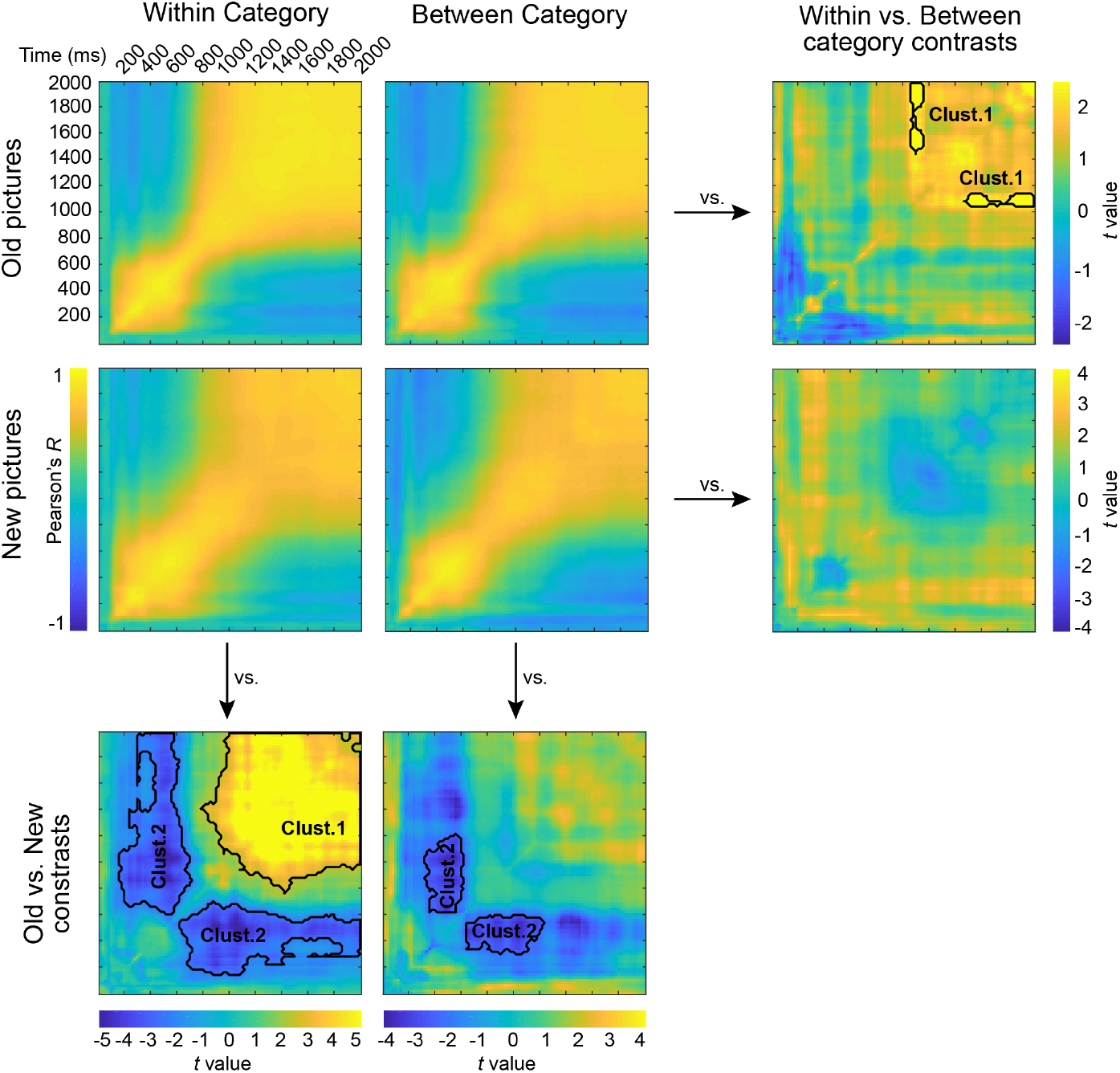
Time-resolved correlation results between trials from the same picture context category (Within condition) and between trials from different context categories (Between condition), separately for EEG responses to Old and New picture cues. Plots depict 2D correlation maps averaged across participants from Experiment 1. Group-level 2D maps of *t*-values are displayed at the edges of the plot depicting results from pair-wise comparisons between conditions. The thick black line depicts the timing of the significant cluster between conditions (p < 0.05, cluster-based permutation test).

Consequently, if observed patterns of stable neural activity were specific to the retrieval of past experiences, then we should expect to find no effects when Within and Between correlations were compared for New responses, as they did not elicit the retrieval of episodic memories. Accordingly, we found no evidence that shifts towards neural stabilization differed between these two correlation outputs in the New responses (Fig. 2). Instead, we found higher stable patterns of activity elicited by New stimuli at early temporal window after stimulus onset (i.e., < 1000 ms). However, this early increase in neural stability to New, versus Old, stimuli was found both when the analysis included trials from the same (Within condition) and from different (Between condition) contextual categories. These findings were further confirmed with a repeated measures ANOVA that revealed a main effect of Response Type (F(1,9) = 6.04, p = 0.03), but not a main effect of Content Type (F(1,9) = 4.54, p = 0.06) nor a Response Type x Content Type interaction effect (F(1,9) = 1.42, p = 0.26). Altogether, the current results suggest, in line with previous electrophysiological studies in humans (Yonelinas et al., 2002; Rugg and Curran, 2007; Staresina and Wimber, 2019), that this early stage of stable activity after stimulus onset may reflect evoked responses associated with novelty detection and familiarity that may precede episodic recollection.

### Retrieval-induced context-dependent EEG stability cannot be attributed to univariate ERP signal properties

We next aimed to further assess whether the distinct temporally evolving patterns of EEG stability associated with New and Old conditions were in fact explained by differences in univariate ERP features that differed between conditions and contexts. The relevance of addressing this issue resides in being able to rule out the possibility they were not reflecting a genuine neural stability process derived from temporally evolving patterns of multivariate neural features but rather by simple differences in scalp ERPs. To address this issue, we compared three different ERP properties: amplitude, topography and variance. First, we contrasted ERP amplitude signal averaged over all scalp electrodes elicited by Old and New picture cues. This analysis revealed that ERPs for Old pictures elicited higher neural response than ERPs for New pictures and that this difference started early on, at ~ 400 ms from cue onset, and remained significantly different for the rest of the window of analysis (Fig. 3a). Similar direction and timing differences were found for ERP variance (Supplementary Fig. 4a) and for scalp topography (Supplementary Fig. 5a). Thus, these findings suggest that Old and New cue pictures elicited different ERP responses that likely explain the correlation differences seen in our previous findings (i.e., Fig. 1f). Importantly, these ERP signal properties did not differ statistically between context conditions for either New or Old cues (Fig 3b; Supplementary Fig. 4b and Supplementary Fig. 5b), thereby excluding the possibility that increases in neural stability at the within level analysis (i.e., Fig. 2) could be mediated by simple univariate ERP signal properties that differ between context picture-types.

**Figure 3.**
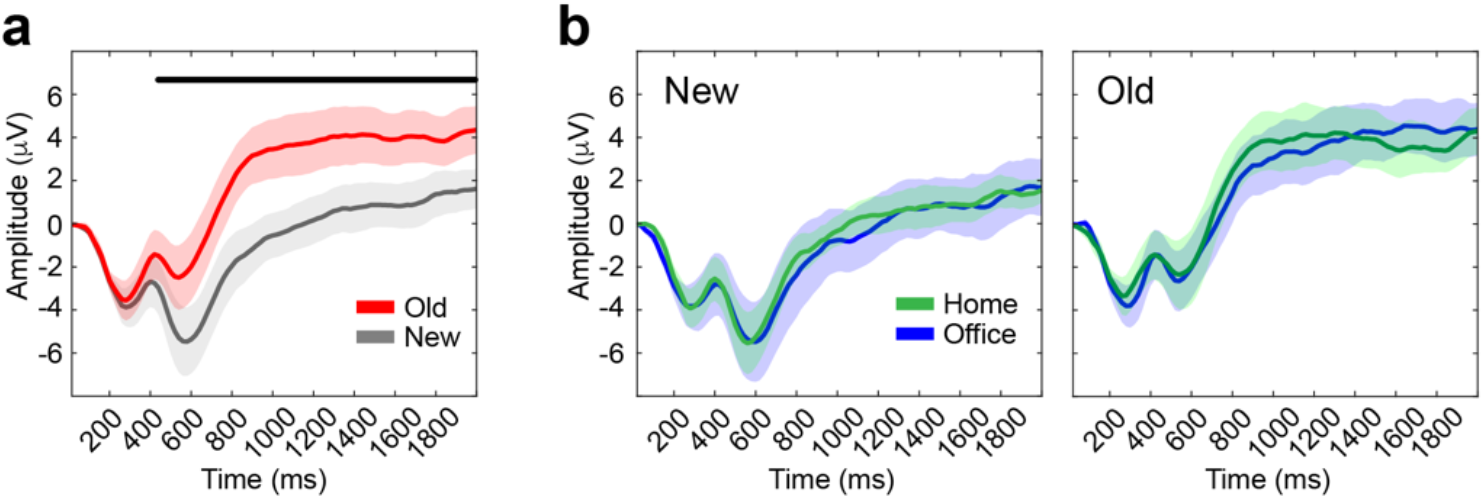
ERPs for picture cues. (**a**) Participants’ averaged ERPs for Old and New pictures presented during the memory recognition task. Thick red and grey lines represent averaged ERPs from all scalp electrodes. Shaded lines around ERPs depict standard error of the mean. Thick black line at top depicts time points at which Old and New ERPs were significantly different (p < 0.05, FDR corrected). (**b**) Similar ERP analysis as in (a) but separating ERPs for pictures displaying image content from “home” and “office” context categories for New and Old conditions.

### Shift toward neural state of stability supports memory recollection

Having shown that stable patterns of activity around ~ 1000 ms from picture onset were associated with memory retrieval, we next sought to investigate whether this effect was associated with participants’ ability to recollect detailed episodic information cued by the picture. To address this issue, we analysed data from a second experiment with a different set of participants *(N* = 11) (see Methods). As in experiment 1, participants from experiment 2 used the wearable camera to record their daily life experiences for a week and were asked to return to the lab to perform a recognition memory test, while EEG was recorded. However, in experiment 2, all pictures presented to the participant during the test were extracted from pictures collected during encoding by the same participant. Thus, participants were not asked to perform an Old/New decision but rather to judge the degree to which each picture elicited ‘Very Low - Low - High’ vivid recollection of the cued past episode. A repeated measures ANOVA, including degree of vividness and picture context category (“home” and “office”) as within-subject factors, confirmed that pictures elicited medium to high degree of vividness to participants and that this was similar for pictures depicting episodes took place at “home” and “at the office” (main effect of degree of vividness: F(2,20) = 8.75, p = 0.002; interaction vividness x category: F(2,20) = 1.49, p = 0.25) (Fig. 4d).

**Figure 4.**
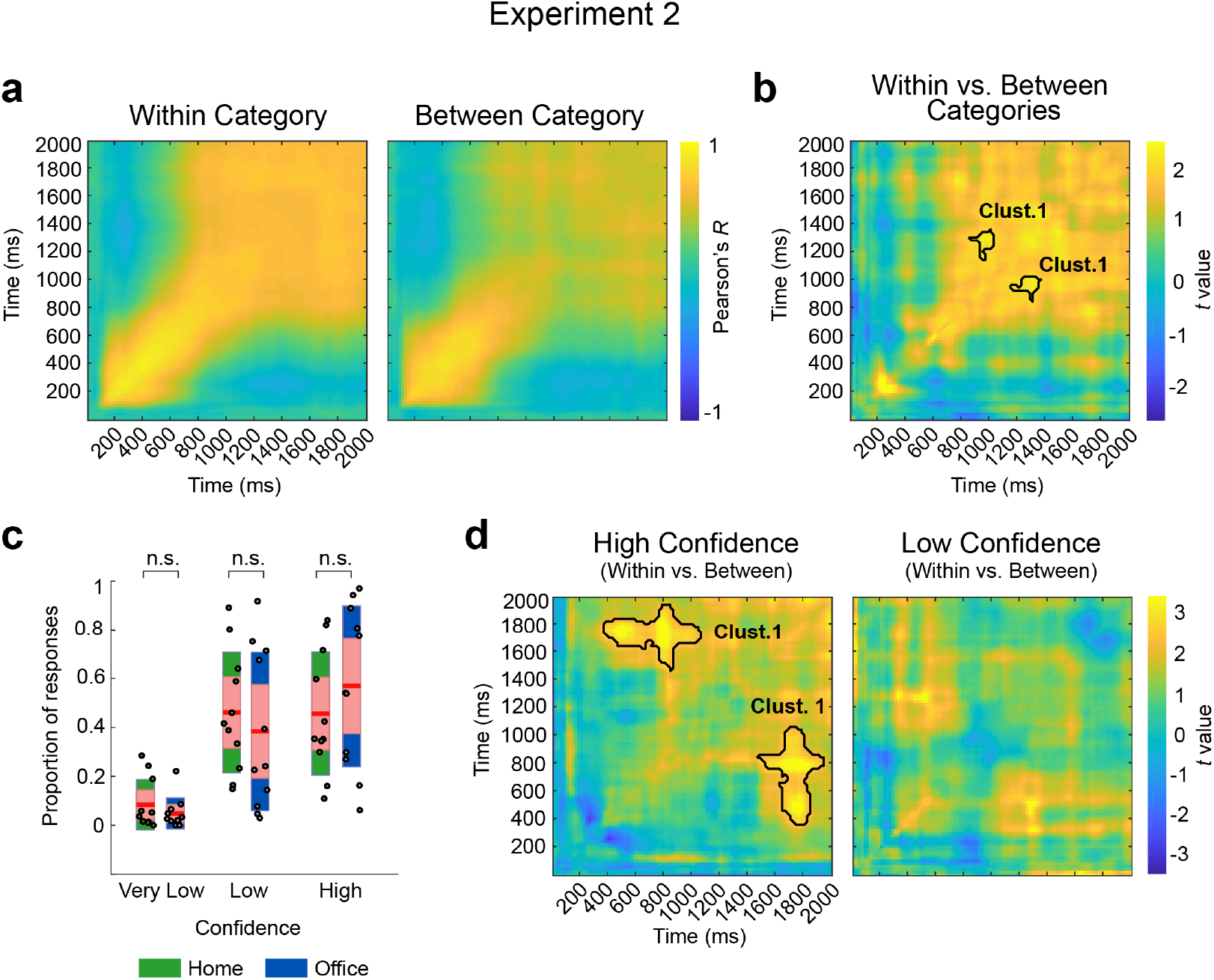
Neural stability and behavioural findings from experiment 2. (**a**) Time-resolved correlation results between trials from the same picture context category (Within condition) and between trials from different context categories (Between condition) for Old picture cues. Plots depict 2D correlation maps averaged across participants. (**b**) Group-level 2D map of *t*-values that results from comparing correlation values between the Within and the Between condition. (**c**) Participants’ proportion of correct responses to Old pictures as a function of the depicted context and confidence judgement. The central mark is the median; the edges of the box are the 25th and 75th percentiles. ‘n.s.’ denotes p > 0.05. (**d**) Group-level 2D map of *t*-values that resulted from comparing correlation values between Within and Between conditions for trials that were judged subjectively by the participants to be High or Low confidence. The thick black line in (b) and (d) depicts the timing of the significant cluster between conditions (p < 0.05, cluster-based permutation test).

First, we examined whether the increase in neural stability found in EEG data elicited by picture cues depicting instances from the same context (Within condition) compared to pictures from different context category (Between condition) also existed in data from experiment 2. Importantly, and corroborating the findings in experiment 1, this finding was replicated in the new dataset. Specifically, we found higher cross-temporal correlation values at ~ 800 ms from picture onset in the Within than in the Between condition analysis (Fig. 4a and b). These findings suggest that shifts towards neural stability can be seen whenever participants are engaged in retrieval tasks, independent of task design.

Then we investigated whether the increase in neural similarity found in the within category condition was functionally relevant to the participants. To address this issue, we organized the recognition trials according to participants’ responses and created two new datasets: one that included EEG data from pictures that triggered vivid recollection of a specific event episode (i.e., those trials followed by “High” responses) and another that included EEG data from pictures that triggered a low degree of episodic vividness (i.e., those trials followed by “Low” responses). Interestingly, only trials in which participants reported vivid picture-elicited episodic recollection showed statistically significant increments in neural stability in the Within as compared to the Between analysis (Fig. 4d). These results indicate the shift towards neural stability is functionally relevant and may support recollection of autobiographical memory episodes.

### Induced neural state pattern of stability is idiosyncratic to each individual

An important question that remains unanswered is whether the retrieval-induced patterns of neural stability are unique to each individual or, alternatively, reflect brain activity states that are common across individuals. Indeed, neuroimaging studies requesting participants to encode and retrieve similar stimuli at the lab have reported that neural patterns of activity elicited at encoding and at retrieval are shared across individuals and that this may reflect common neural representations for shared knowledge (Chen et al., 2017). In the context of retrieving real-life autobiographical episodic memories, however, the notion of shared representations is complicated, as one’s own past experience is genuinely individual. To examine this issue, we implemented a cross-temporal Inter-Subject Pattern Similarity (ISPS) analysis of the data from experiment 1, which allowed us to examine how well temporally evolving EEG patterns of stability elicited by Old and New pictures were shared across individuals (see Methods). In line with the notion that neural stability patterns were idiosyncratic to each individual during the retrieval of real-life autobiographical memories, we found that an increase in correlation values in the Within compared to the Between condition was no longer observable (Fig. 5). However, the two task-oriented and temporally defined clusters of neural stability associated with Old and New responses found at the individual level were also found in this across-subject analytical approach. More concretely, we found higher ISPS values associated to the presentation of New stimuli at an early temporal window (i.e., < ~1000 ms after stimulus onset) and higher ISPS values associated to the presentation of Old stimuli at a later temporal window (i.e., > ~1000 ms after stimulus onset). However, this pattern of results was found when the across-subjects correlation approach included EEG data elicited by pictures from the same (Within condition) and from different context category (Between condition) (Fig. 5). A repeated measures ANOVA corroborated the increase neural similarity in cluster 1 and2 was driven by a difference between Old and New conditions rather than differences in the Within and Between type of analysis (Cluster 1: significant main Response Type effect (F(1,9) = 16.64, p = 0.003) but not a Content type (F(1,9) = 4.03, p = 0.07) or an interaction effect (F(1,9) = 1.20, p = 0.30); Cluster 2: significant main Response Type effect (F(1,9) = 33.02, p < 0.001) but not a Content type (F(1,9) = 2.71, p = 0.13) or an interaction effect (F(1,9) = 2.64, p = 0.14)). Thus, while these findings suggest that individuals share general neural response pattern dynamics elicited by the identification of Old or New incoming cues, they also indicate that shifts towards stable state patterns of activity during retrieval are idiosyncratic to each individual and associated with the retrieval of their own real-life autobiographical memories.

**Figure 5.**
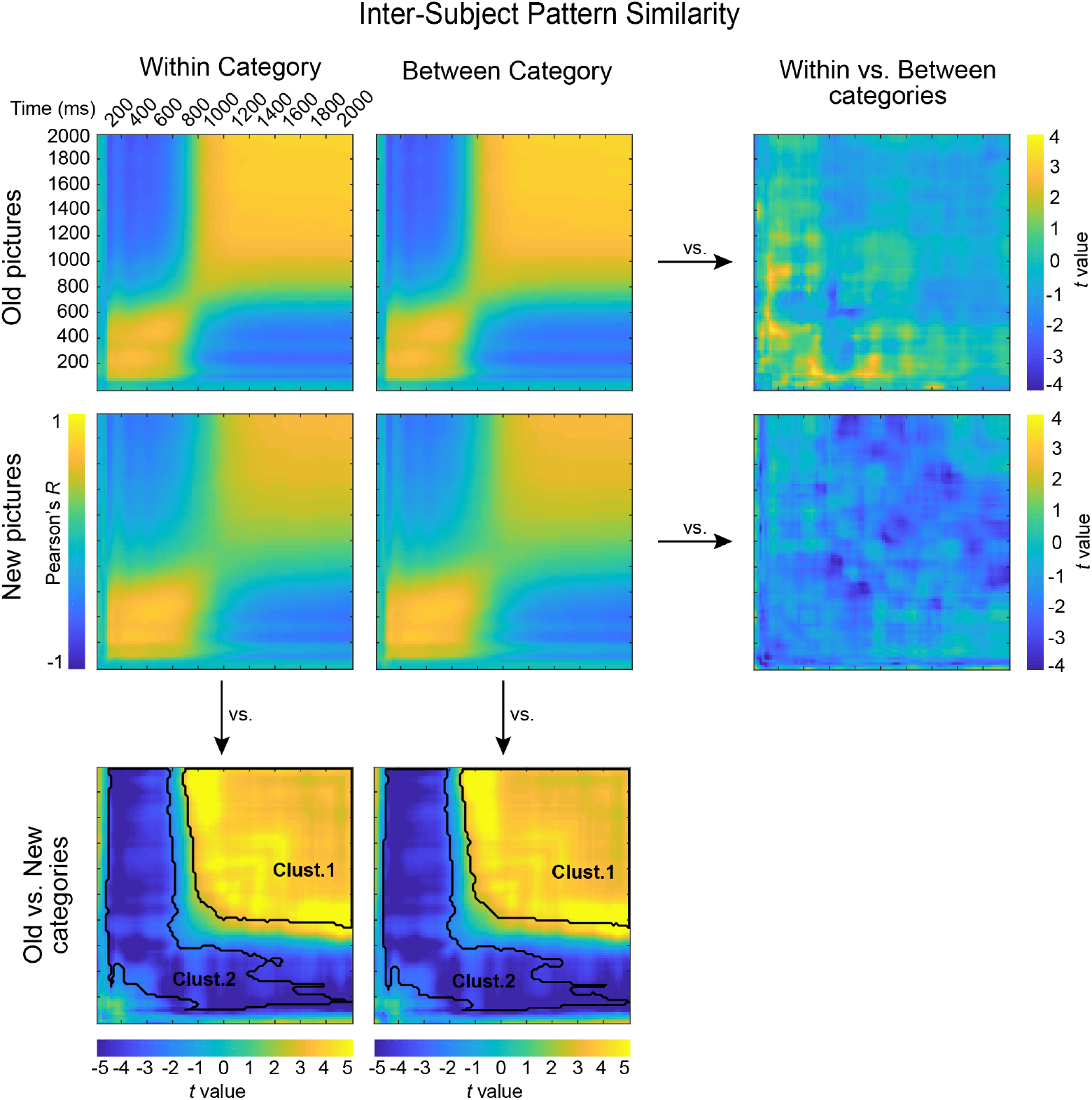
Inter-subject pattern similarity in experiment 1. Inter-subject correlation value derived from correlating the patterns for each experimental condition in each individual with the corresponding EEG patterns’ condition in the rest of the group. Group-level 2D maps of *t*-values are displayed at the edges of the plot depicting results from pair-wise comparisons between conditions. The thick black line depicts the timing of the significant cluster between conditions (p < 0.05, cluster-based permutation test).

## Discussion

The current study offers three novel findings. It shows that the retrieval of real-life episodic memories undergoes a rapid shift towards brain patterns of stable activity, that the degree of neural stability is associated with participants’ ability to recollect the content of the cued episode, and that the stabilized neural pattern during retrieval is idiosyncratic to an individual and not shared across other participants. These results indicate that the ability to retrieve autobiographical memory episodes is supported by rapid shifts of neural activity towards stable states.

The notion that a subtle reminder can bring back memory episodes from the past is arguably one of the main pillars of cognition and behaviour. Ignited by neuropsychological work and corroborated by animal models and human neuroimaging converging evidence points to the medial temporal lobe (MTL), and the hippocampus in particular, as the key brain region supporting episodic memory (Davachi, 2006; Scoville and Milner, 1957; Squire, 1992). Based on the physiological properties of the hippocampal CA3 subregion, computational models have proposed that the dense recurrent connections among CA3 pyramidal cells engender auto-associative network processes characterized by attractor-like dynamics that render activity towards a stable state (Marr, 1971; Rolls, 2016). The stable activity states they produce are called ‘attractors’, because when a stimulus causes the network’s activity to become similar to, or ‘nearby’ the attractor, the internal dynamics of the network cause that activity to shift towards (or be ‘attracted to’) the attractor state. As a result, the hippocampal circuitry itself enables the so-called ‘pattern completion’ and subsequent reinstatement at the neocortex (McClelland et al., 1995; Treves and Rolls, 1992; Norman and O’Reilly, 2003). Our finding that only picture cues depicting past episodes elicited stable states of neural activity from ~ 800 ms after input onset illustrates the temporal dynamics that account for attractor-like patterns of activity during episodic retrieval, an important characteristic of this mechanism that had not been revealed previously.

Though pattern completion has been central to influential theoretical models of memory retrieval (Marr, 1971; McClelland et al., 1995; Rolls, 2016), it is only recently that we have received evidence for its existence in humans (Staresina et al., 2016; Horner et al., 2014 and 2015; Treder et al., 2020). This research has been successful in identifying neural signatures underlying pattern completion by using lab-based stimulus material thought to emulate episodic memory functioning in real-life. The current findings extend these results by showing that shifts towards stable neural activity occur when individuals retrieve autobiographical events encoded during everyday life activity, thereby lending support to the notion that this mechanistic memory principle mediates memory retrieval in real-life.

Our findings also show that picture cues collected by each participant depicting episodes from the same context, i.e., at home or at the office during everyday life routine, elicit stable neural activity dynamics not shared across participants, thereby supporting the theoretical prediction that stable states of neural activity are content-specific in an individual’s memory (Treves and Rolls, 1992). Recent findings using ISPS analysis on neuroimaging data recorded while people watched a movie demonstrated the existence of a common spatial organization in high-order brain regions for memory representations across participants (Chen et al., 2017). The observation that neural patterns of activity during memory encoding and retrieval are similar across individuals has been replicated in several other studies using neuroimaging (Baldassano et al., 2017; Raykov et al., 2020) and scalp electrophysiological (Silva et al., 2019) recording techniques, thereby suggesting that people share similar abstract representational codes when encoding and retrieving similar stimuli, even if it is ecologically valid like a movie. However, memories of real-life events are fundamentally unique and dependent on each person’s experience, thereby precluding the possibility that memory representations are shared with others. Thus, our findings emphasize the idiosyncratic nature of memory representations derived from real life, and they are congruent with general theories of memory organization (i.e., Multiple Trace Theory; Nadel and Moscovitch, 1997) and autobiographical memory models (Conway and Pleydell-Pearce, 2000) that emphasize the strong link between abstract memory representations and representations of individual memory episodes. Arguably, an interesting hypothesis for future work would be to assess whether shared neural patterns elicited by picture cues from a context are shared by individuals who share similar real-life experiences, such as those who live in the same place or who work in the same office. An additional intriguing possibility would be to examine whether shared patterns of neural stability elicited by real-life cues depicting same-context representations, as in our study, are mediated by social network proximity parameters such as friendship (e.g., Parkinson et al., 2018).

In summary, our findings reveal the existence of rapid shifts towards stable activity patterns during episodic memory retrieval. At the same time, they highlight the importance of studying temporally evolving neural signals and how attractor-like dynamics emerge rapidly upon partial cues of an experienced past episode. Taken together, these findings help bridge the fields of theoretical and empirical neuroscience in learning and memory and shed light on neural mechanisms that promote memory recollection of real-life autobiographical episodic memories.

## Methods

### Design overview

The study included data from two separate experiments on two different samples of healthy participants. In the two experiments, participants were instructed to carry a wearable camera during their routine daily life activity over a period of 5-7 consecutive days. Participants were asked to come to a training session a few days before the start of the study. In this session they were informed about how the camera worked, and they read, understood, and signed the informed consent. We took special care in providing details on privacy issues so that all participants were fully aware of them before the beginning of the study. The camera took face-front pictures automatically every 30 seconds and participants reported their activity was not altered because of this during the week. Thus, most of the participants’ collection of pictures depicted images from episodes taking place at “home” or at “the office”, though other everyday life activities were also reflected, such as being “on public transport” or “in the street”. Participants were asked to return the materials to the experimenter for processing at the end of the encoding week. Participants confirmed not having checked the pictures during the encoding week. In experiment 1, a recognition memory test was administered one and two weeks after encoding. Pictures depicting episodes in the participant’s past were presented only once in the two tests and their presentation was interspersed with pictures from another participant’s past, which allowed assessment of participants’ ability to distinguish Old from New information (Fig. 1a-c). In experiment 2, participants’ memory was assessed via a recognition memory test administered one week after encoding. All pictures presented during this test were extracted from each participant’s collected set of acquired pictures and they were requested to indicate the extent to which each of the picture cues elicited a vivid memory from their past.

### Participants

In experiment 1, fourteen healthy (eight women), right-handed, adults with normal or corrected vision participated in the study. The age range was 22-37 years old (average age = 27.68 years old; SD = 4.22). However, EEG data from two participants were lost due to technical problems and were not able to be included in the rest of the analysis. In experiment 2, sixteen (eight women, mean age = 26.7) healthy, right-handed, adults with normal or corrected vision were recruited. None of the participants suffered from any neuropsychiatric disorders. All participants provided informed written consent for the protocol approved by the Ethics Committee of the University of Barcelona. Participants received financial compensation of 60€ and 40€ for their participation in each of the experiments, respectively.

### Wearable camera

We used the wearable Narrative clip 2 camera (http://getnarrative.com/) with an 8MP sensor that can be attached to the person’s clothing or can be carried as a necklace using a thin cord. The camera’s lens includes a light sensor which is automatically triggered by the light intensity. This provides automatic capture of high-resolution photographs (3264 x 2448 px) every thirty seconds. The camera does not include a digital screen, which prevents the participants from having direct access to the captured pictures. However, each camera was provided with the company’s uploading software (Narrative Uploader), through which the participants were able to locally store each day’s pictures on their personal computer. At the end of the recording week, all photographs were delivered to us through a USB stick. The camera was programmed to take images automatically every thirty seconds and produced pictures with an egocentric point of view.

### Picture selection

In experiment 1, we implemented a deep neural networkbased algorithm, SR-Clustering (Dimiccoli et al., 2015), to automatically organize the stream of each participant’s pictures into a set of temporally evolving meaningful events. The algorithm segments picture sequences into discrete events (event segmentation; e.g., having breakfast in a kitchen, commuting to work, being in a meeting) on the basis of its ability to identify similar contextual and semantic features from the picture stream. The implementation of the SR-Clustering algorithm provided a variable number of discrete events for each participant per day, and each event included 8 to 20 pictures. Each participant’s events were then manually inspected and those which displayed non-meaningful episodes (e.g., pictures were blurred, or the camera was pointing to the roof or was blocked by clothing) were discarded. Picture events that represented interactions with participants’ relatives were excluded from the study. Three independent experimenters rated and selected the set of event pictures for each participant on the basis of these criteria, and only those events that were consistently selected by the three investigators were included in the final set of picture events in the study. Once the images were organized into discrete events, we selected a representative picture from each event, thereby ensuring most of the past episodic experience was brought into the test. We then numbered the sequence of event pictures and assigned even-numbered pictures to be used as memory cues for the first test (one week after the encoding week) and odd-numbered pictures to be included in the second test (two weeks after the encoding week). Picture cues presented to one participant depicting their own past (Old) were also presented to another participant as New images. This ensured that differences between Old and New pictures presented to each participant were only based on the image’s direct link to ones’ past while preserving the rest of the picture characteristics intact during the test (e.g., angle of view, description of routine daily life activities).

In experiment 2, ~ 80 pictures were carefully chosen to represent each participant’s daily activity. This added up to a total number of ~400 stimuli used in the recognition memory test. The selection of the pictures was made manually by two separate experimenters following the criterion that each picture should correspond to a single episodic event (defined by a sequence of pictures depicting instances in the same spatiotemporal context). If number of event pictures exceeded 80 per day for each of the two experimenters, pictures depicting episodic instances with less clear beginning and ending moments in time were discarded.

None of the participants in the two experiments had personal relationships with each other, nor did we encounter an instance where two concurrently enrolled participants came into direct contact with one another while wearing their cameras.

### Recognition memory task

In experiment 1, participants were asked to distinguish pictures that depicted instances from their own past (Old condition) from those that did not (New condition). Pictures were presented on the screen for 3000 ms with a message “Old or New?” prompting participants to indicate on a keyboard whether the picture displayed experienced (Old) or novel (New) events. Old and New pictures were presented in a random order during the task. Old and New pictures were presented in a random order during the task. Participants were then asked to judge whether they were “Sure/Not sure” when they indicated an image was “New” and “Remember/Know/Guess” when images were perceived as “Old”. They were instructed that “Guess” meant they had no contextual memory reference for what was depicted in the image, but they recognized the content as being from their own life (e.g., viewing ones’ living room). “Know” was to be selected when the visual content in the picture was highly familiar but the subject could not determine what took place in it, perhaps because the event was part of a routine (e.g., playing football on Thursdays), while “Remember” related to when the picture elicited a vivid memory of that specific event and could be located in time. Participants’ ability to order each event depicted in the pictures during the encoding week was tested afterwards more concretely, when they were asked to indicate whether the pictures depicted an event that took place at the “beginning, middle, or end” of the encoding week as the image appeared on the screen. Finally, participants were asked to rate the degree to which each of the pictures elicited an emotional response and to indicate whether it was positive or negative on a scale that ranged from 2 (maximum positive) to −2 (maximum negative), with 0 being neutral emotionally. Details of participants’ confidence judgement, temporal order, and emotional ratings can be found elsewhere (Nicolás et al., 2020). Briefly, most of participants stated that they were “sure” and “know” when indicating New and Old responses, respectively. In general, they showed difficulties in temporally ordering the episodic events depicted by Old pictures during the test, and they reported neutral emotional ratings to most of the pictures. This pattern of behavioural results was similar in the two tests of the experiment.

In experiment 2, all pictures presented were extracted from those collected from each participant’s encoding week. The order of picture presentation preserved the temporal order from encoding. Thus, the recognition memory test recreated the sequence of event episodes experienced in real-life. The structure of each trial was as follows: it started with the appearance of a fixation cross for a duration of 1000 ms in the centre of the screen, followed by display of the picture image for 3000 ms. A confidence judgement task followed. A message on the screen prompted participants to indicate they had been able to retrieve episodic details associated with the just-presented picture cue with “Very low/Low/High” confidence.

### EEG recording and preprocessing

EEG was recorded (band-pass filter: 0.01–250 Hz, notch filter a 50 Hz, and 500 Hz sampling rate) from the scalp using a BrainAmp amplifier and tin electrodes mounted in an electrocap (Electro-Cap International) located at 29 standard positions (Fp1/2, Fz, F7/8, F3/4, FCz, FC1/2, FC5/6, Cz, C3/4, T3/4, Cp1/2, Cp5/6, Pz, P3/4, T5/6, PO1/2, Oz) and at the left and right mastoids. An electrode placed at the lateral outer canthus of the right eye served as an online reference. EEG was re-referenced offline to the linked mastoids. Vertical eye movements were monitored with an electrode at the infraorbital ridge of the right eye (EOG channel).

In the current study (both experiment 1 and 2), only EEG data elicited by picture cues depicting episodic instances taking place at participants’ “home” and “office” were analysed.

### Event-related potential (ERP) analysis

The continuous sample EEG data were then epoched into 2100 ms segments (0 to 2000 ms relative to trial onset), and the pre-stimulus interval (−100 to 0 ms) was used as the baseline for baseline correction procedure. Trials exceeding ± 100 μV in EEG and/or EOG channels within a −100 to 2000 ms time window from stimulus onset were rejected offline and not used in ERPs and time-frequency analysis (see details below). For each participant, we obtained trial epochs that were separately catalogued as belonging to Old and New conditions.

### EEG pattern stability

First, to account for whether Old and New pictures elicited different patterns of neural stability, a temporally resolved autocorrelation analysis was implemented at single trial level using Pearson correlation coefficients (*R*), which are insensitive to the absolute amplitude and variance of the EEG response. The autocorrelation analysis on EEG data elicited by picture cues was made at the individual level, and included spatial (i.e., scalp voltages from all the 29 electrodes) and temporal features, which were selected in steps of 20 sample points (40 ms) of the resulting *z*-transformed EEG single trials. The resulting correlation matrices for Old and New trials were then averaged separately, which produced an individual 2D matrix of time-resolved degree of pattern stability (Fig. 1d).

To examine whether context-specific information depicted in picture cues (i.e., “home” or “office”) modulated the degree of EEG response stability we randomly assigned an individual’s *z*-transformed EEG data from the same category into two sets of equal number trials. A time-resolved correlation analysis similar to that described above was then implemented on EEG data resulting from averaging the two sets of EEG trials separately. This approach allowed random selection of the averaged EEG data from one of the two sets from category A and correlation of it with the averaged EEG data from the other set, resulting in a within category correlation matrix, and with one of the two sets from category B, resulting in a between-category correlation matrix.

In the current study, participants that did not have at least 30 trials per picture context category (“home” and “office”) to each experimental condition (Old and New) in experiment 1 and 16 trials to each context category to each degree of vividness in experiment 2 were ruled out from the analysis. As a result, data from one participant from experiment 1 and five from experiment 2 were not included in the analysis.

### Inter-subject pattern similarity analysis (ISPS)

We examined whether neural stability patterns elicited by picture cues from the same category (i.e., within) and from different categories (i.e., between) were similar across participants in experiment 1. To address this issue, we computed a similar time-resolved correlation analysis between averaged EEG neural response trials for each category from one participant with the same (i.e., within condition) or different (i.e., between condition) averaged EEG neural response trials averaged across the remaining participants, for each left-out participant. The average of the resulting matrices resulted in an across-participants similarity analysis.

### Cluster statistics on EEG data

To account for neural similarity differences between conditions a cluster-based permutation test was used (Maris and Oostenvelt, 2007). This approach searches for clusters of significant points in the resulting spatiotemporal 2D similarity matrix (time x time) in a data-driven manner and addresses the multiple-comparison problem by employing a nonparametric statistical method based on cluster-level randomization testing to control for the family-wise error rate. Statistics were computed for each time point, and the temporal points whose statistical values were larger than a threshold (p < 0.05, two-tail) were selected and clustered into connected sets on the basis of x, y adjacency in the 2D matrix (defined by at least 2 contiguous points). The observed cluster-level statistics were calculated by taking the sum of the statistical values within a cluster. Then condition labels were permuted 1000 times to simulate the null hypothesis, and the maximum cluster statistic was chosen to construct a distribution of the cluster-level statistics under the null hypothesis. The nonparametric statistical test was made by calculating the proportion of randomized test statistics that exceeded the observed cluster-level statistics.

Differences in ERP amplitude between conditions were assessed via paired-sample Student t-test. We used a paired sample permutation test (Groppe et al., 2011) to deal with the multiple comparisons problem given the number of sample points included in the analysis. This test uses the “tmax” method for adjusting the p-values of each variable for multiple comparisons (Blair and Karniski, 1993). Like Bonferroni correction, this method adjusts p-values in a way that controls for the family-wise error rate.

## Acknowledgments

We thank Valeria Marzano for assistance in data collection, Àngel Bujalance in data preparation and Albert Compte, Klaus Wimmer, Rubén Moreno-Bote and Josep Marco-Pallarés for their helpful discussions on earlier versions of the manuscript. This work was supported by Ministerio de Ciencia, Innovación y Universidades, which is part of Agencia Estatal de Investigación (AEI), through the project PSI2016-80489-P (Co-funded by European Regional Development Fund. ERDF, a way to build Europe), to L.F. The project that gave rise to these results received the support of a fellowship from “la Caixa” Foundation (ID 100010434). The fellowship code is LCF/BQ/DI19/11730040. We thank CERCA Programme/Generalitat de Catalunya for institutional support.

## Author Contributions

L.F., B.N. and A.K. conceived the research. L.F., B.N. and A.K. designed the research. B.N. and A.K. performed the research. L.F. and M.S. analyzed data. L.F., B.N. and A.K. and M.S. wrote the manuscript. All authors discussed the results and contributed to the manuscript.

## Supplementary Information

**Supplementary Figure 1.**
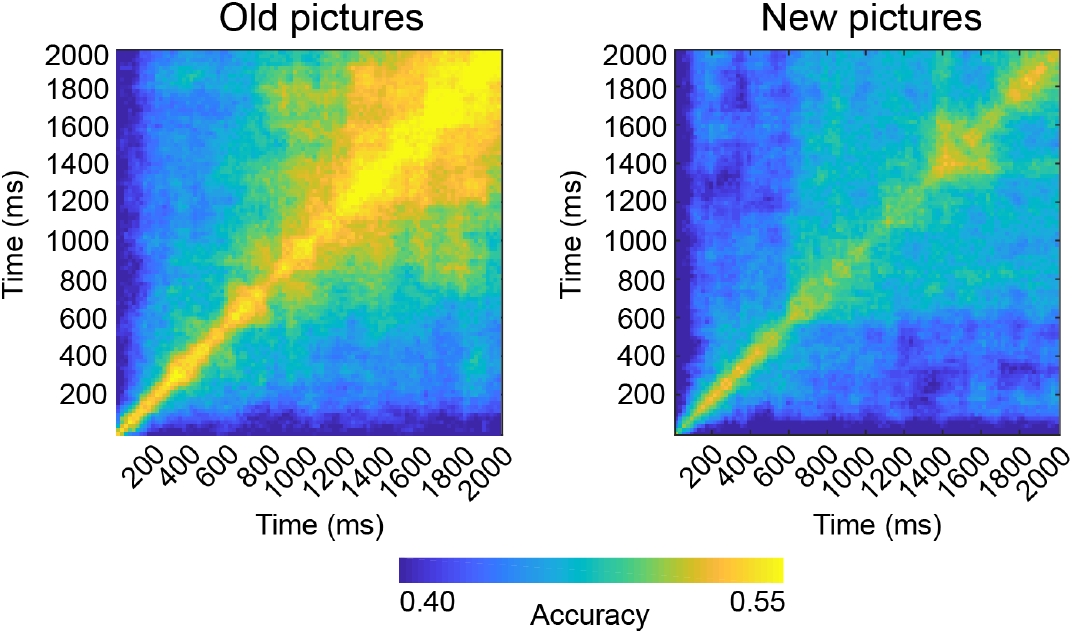
Linear discriminant analysis (LDA) results in experiment 1. An LDA was trained and tested on the EEG sensor patterns (pre-processed signal amplitude on each of the 29 channels), independently per participant and cue condition (Old and New) and at each sample point during retrieval from picture onset (0 ms) to 2000 ms post-cue. The classifier was trained to detect systematic differences between trials of pictures depicting “home” and “office” content. A leave-one-out cross-validation procedure was used to train and test the classifier. Time resolved LDA results for EEG to Old and New pictures were averaged and displayed in this figure. An increase in LDA accuracy can be observed at 1000 ms from cue onset for Old pictures. The timing of the LDA increases coincides with increased correlation findings described in the study for Old pictures.

**Supplementary Figure 2.**
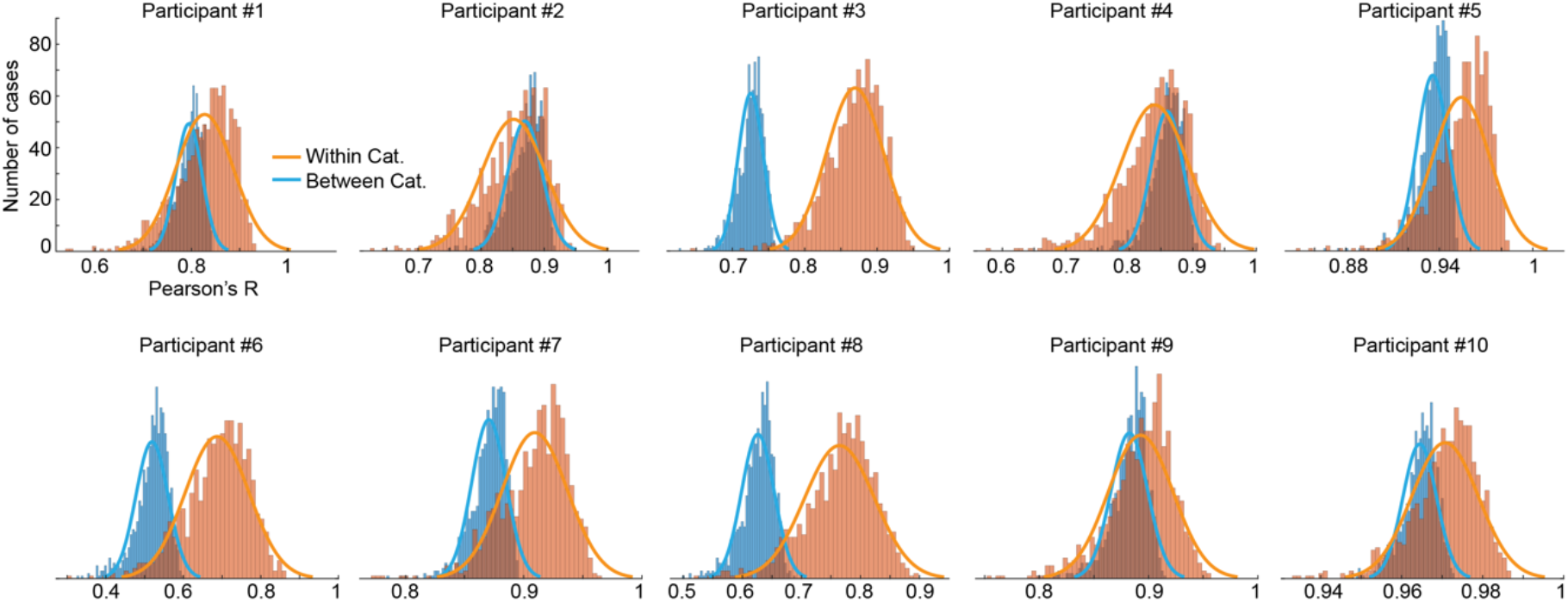
Distribution of Pearson *R* values to permutation analysis run at individual level in experiment 1. To exclude the possibility that, in experiment 1, the increase in neural stability found in the main analysis in the within compared to the between condition was artificially induced by how EEG trials were split into two sets before running the correlation analysis, we created, for each individual, a set of 1000 random trial assignments and implemented the neural similarity analysis accordingly. This figure displays the distribution of Pearson *R* values for each participant. We then averaged each individual’s 1000 correlation values obtained for each of the conditions and compared them at the group level by means of a paired t-test. This analysis corroborated previous findings for correlation values in the cluster remained significantly higher in the Within than in the Between condition in the Old picture condition (t(9) = 2.32, p = 0.045).

**Supplementary Figure 3.**
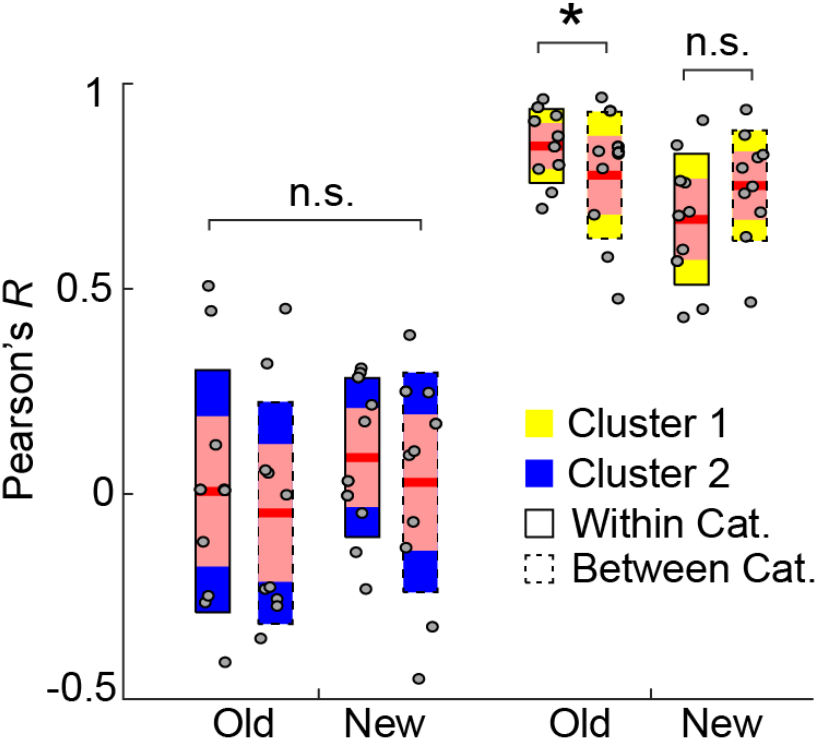
Correlation values associated with each experimental condition in experiment 1. Individual correlation values were extracted from the average for all sample points included in significant cluster 1 found in the group level analysis displayed in Fig. 2 in the manuscript. For all boxplots, the central mark is the median, and the edges of the box are the 25th and 75th percentiles. ‘n.s’ denotes p > 0.05 and * denotes p < 0.05.

**Supplementary Figure 4.**
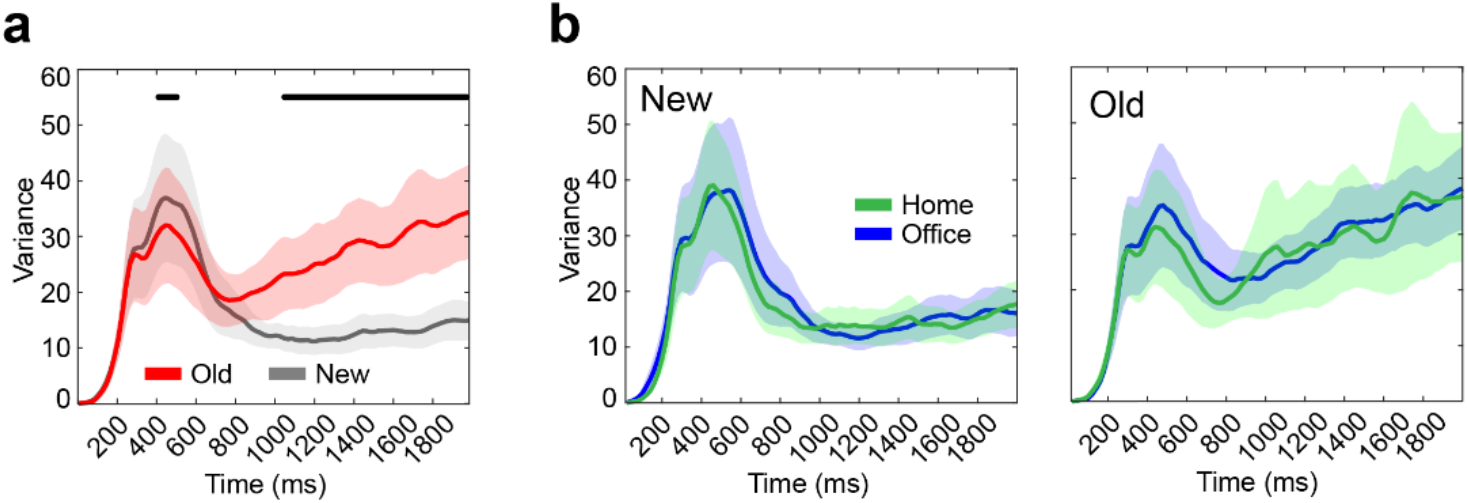
Time resolved ERP variance results in experiment 1. (**a**) Participants’ averaged variance across Old and New pictures presented during the memory recognition task. Thick red and grey lines represent averaged ERPs from all scalp electrodes. Shaded lines around ERPs depict standard error of the mean. Thick black line at the top depicts time points at which Old and New ERPs were significantly different (p < 0.05, FDR corrected). (**b**) Similar ERP analysis as in (a) but separating ERPs into pictures displaying image content from “home” and “office” context categories for New and Old conditions.

**Supplementary Figure 5.**
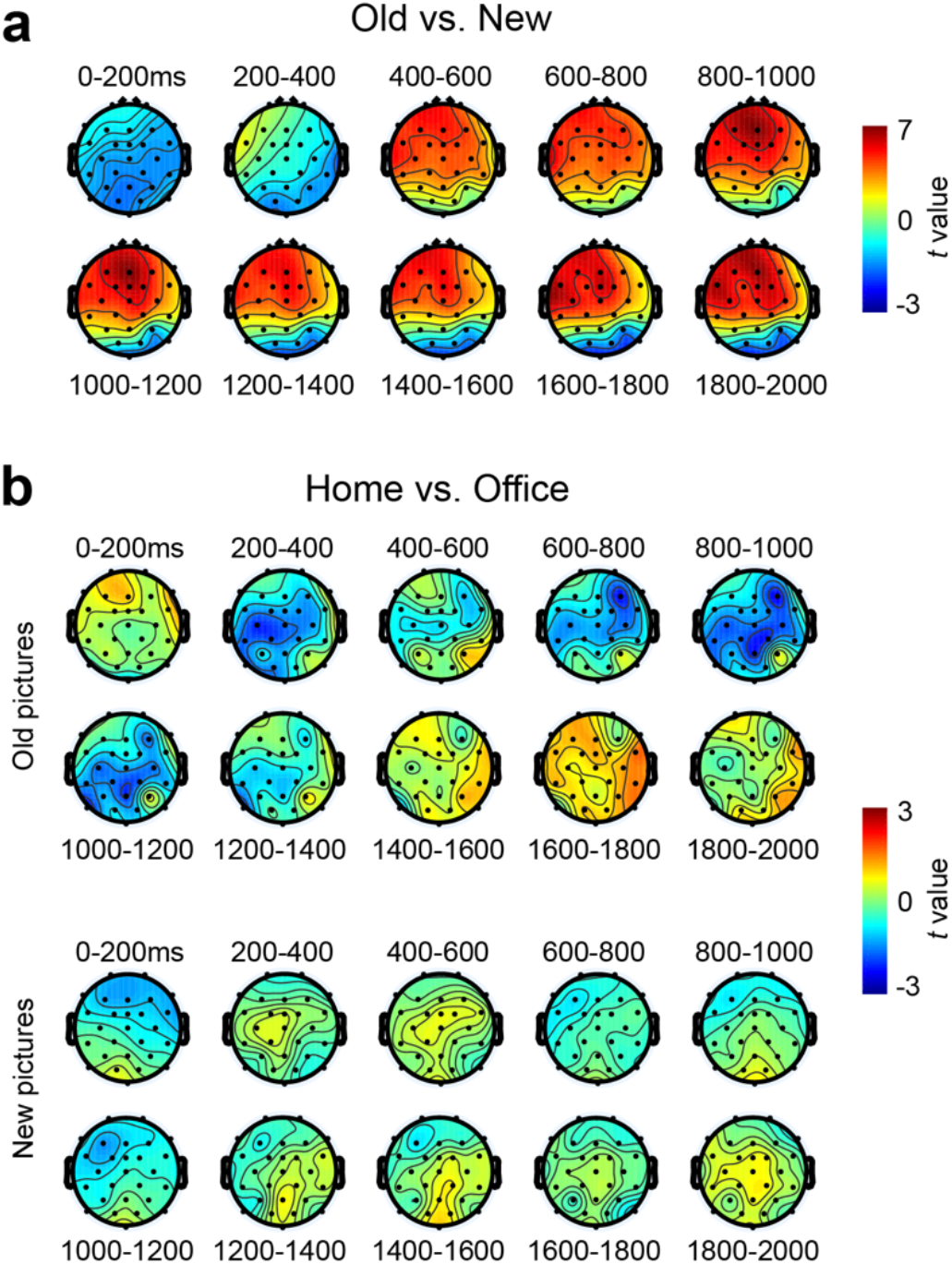
(**a**) Scalp topography of ERP amplitude differences elicited by Old and New pictures. Topography maps depict electrode-by-electrode t-values (paired Student t-test) that resulted from comparing participants’ ERP signal averaged over windows of 200 ms from 0 to 2000 ms at picture onset. (**b**) Scalp topography of ERP amplitude differences (expressed in t-values) elicited by Home and Office picture cues separately for Old and New conditions.

